# Obesity Increases Atherosclerosis Susceptibility via Inter-tissue miR-30e-SLC7A11 Axis

**DOI:** 10.1101/2025.03.06.641950

**Authors:** Chen Wang, Jianqing She, Xiaoli Qian, Haoyu Wu, Wenyang Hao, Xiao Liang, Ning Guo, Xulei Dai, Fangzhou He, Songyang He, Jin Zhang, Yuyang Lei, Xiaozhen Zhuo, Hongbing Li, Yongbai Luo, Kai Deng, Yi Liu, Shanshan Gao, Xiao Yuan, Hui Liu, Ting Bai, Ying Xiong, John Y-J. Shyy, Zu-Yi Yuan

**Affiliations:** Department of Cardiology, The First Affiliated Hospital of Xi’an Jiaotong University, Xi’an, China; Cardiovascular Research Center, School of Basic Medical Sciences, Xi’an Jiaotong University Health Science Center, Xi’an, China; Division of Cardiology, Department of Medicine, University of California, San Diego, La Jolla, CA; Department of Laboratory Animal Science, School of Basic Medical Sciences, Xi’an Jiaotong University, Xi’an, China; Department of Clinical Laboratory, The First Affiliated Hospital of Xi’an Jiaotong University, Xi’an, China; BioBank, The First Affiliated Hospital of Xi’an Jiaotong University, Xi’an, China

**Keywords:** Obesity, MiR-30e, Endothelial cell, SLC7A11, Atherosclerosis

## Abstract

**Background:** With obesity as a risk factor for atherosclerotic disease, recent research suggests that adipose tissue in obese animal models and humans can generate endocrine-like molecules that affect arterial health later on. Previous studies showed that microRNA-30e (miR-30e) level is elevated in atherosclerosis and solute carrier family 7 member 11 (SLC7A11), a cystine/glutamate transporter, is involved in atherogenesis. However, whether an endocrine-like link between the adipose-derived miR-30e-5p and SLC7A11 in the vascular endothelium can lead to obesity-caused atherosclerosis is unknown.

**Methods:** MiRNA data mining and RT-PCR validations were used to demonstrate the positive association among serum level of miR-30e-5p, obesity, and coronary arterial disease in human patients. Transcriptomics (RNA-seq and single-nucleus RNA-seq), metabolomics, and *in silico* analysis were used to establish a miR-30e-5p–SLC7A11 regulation of central carbon metabolism, mitochondrial and endothelial cell (EC) function. Mouse models with EC-specific *Slc7a11* knockout (EC-*Slc7a11*^-/-^) and gain or loss of function of miR-30e-5p were used to elucidate the detrimental role of this endocrine-like axis in obesity-related atherosclerosis.

**Results:** The level of adipocyte-derived miR-30e-5p was significantly upregulated in obese humans with coronary artery disease and obese and atherosclerotic mice. Via serum exosomes, the adipocyte-generated miR-30e-5p targeted *SLC7A11* mRNA in vascular ECs. SLC7A11 deficiency due to miR-30e-5p targeting dysregulated glutamate/cystine metabolism increased glycolysis, reduced oxidative phosphorylation, and impaired mitochondrial function. The EC dysfunction could be rectified by SLC7A11 overexpression or miR-30e-5p antagonism. Exogenously delivered miR-30e-5p phenocopied the increased atherosclerosis in EC-*Slc7a11*^-/-^ mice. In contrast, miR-30e-5p antagomir treatment reduced atherosclerosis in *Apoe*^-/-^ and *ob/ob* mice.

**Conclusion:** Our multi-omics approaches demonstrates that the adipose-derived miR-30e-5p downregulated *SLC7A11* mRNA in ECs via tissue crosstalk. The resulting EC dysfunction led to obesity-related atherosclerosis in mice. These findings underscore a causality between obesity and atherosclerosis in the context of cardiovascular-kidney-metabolic syndrome.

## Introduction

Atherosclerosis is a multi-faceted disease that involves chronic inflammation and maladapted metabolism in several tissue types.^1–3^ Clinical guidance from the National Cholesterol Education Program Adult Treatment Panel III (NCEP ATP III) and World Health Organization suggest that patients with metabolic syndrome are at greatly increased risk of atherosclerotic cardiovascular disease.^4,5^ The novel concept of cardiovascular-kidney-metabolic (CKM) syndrome has brought further attention to the causation between obesity and atherosclerosis.^6^ Despite ample clinical evidence revealing the association between metabolically impaired adipose tissue and atherosclerotic vasculature, the underlying mechanism remains unclear.

Knowledge gained from multiple studies shows a strong association between visceral adipose tissue and atherosclerotic outcomes.^7,8^ At the onset of obesity, the impaired differentiation of adipocytes reduces the production of adiponectin but increases the secretion of proinflammatory cytokines such as tumor necrosis factor α (TNF-α), interleukin 6 (IL-6), and monocyte chemoattractant protein-1.^9^ The prevalent hypothesis of atherosclerosis states that endothelial injury aggravates vascular inflammation and lipid uptake, which initiates atherosclerosis.^10,11^ Extracellular vesicles, including exosomes, play a crucial role in intercellular communication by transferring DNA, coding and noncoding RNA, and proteins to the extracellular space as well as tissues nearby and far.^12^ Because adipocytes are an abundant tissue origin of exosome-carried microRNAs (miRNAs) in the circulation, emerging evidence suggests that adipocyte-derived exosomes can regulate vascular function and disease in an “endocrine-like” manner.^12,13^ However, whether any exosome-associated miRNA can function as a messenger between adipose tissue and vascular endothelium is unknown, as is the effect on atherosclerotic cardiovascular disease.

The p53-induced miR-30e has been recognized as a tumor suppressor in multiple cancer types, including hepatocellular carcinoma, colorectal cancer, and breast cancer.^14–16^ New evidence reveals that miR-30e is linked to atherosclerotic cardiovascular disease (ASCVD) in that circulating miR-30 levels are associated with coronary calcification and plaque instability and elevated level of miR-30e is evident in atherosclerotic lesions.^17–19^ In terms of vascular endothelial cells (ECs), circulating extracellular vesicles (EVs) from diabetic (*db/db*) mice show an increased amount of miR-30e-5p, which coincides with coronary microvascular EC dysfunction and diastolic dysfunction.^20^ Of note, miR-30e is primarily released from adipose tissue in an exosomes-dependent manner.^12^ However, how adipose-derived miR-30e affects the endothelium to exacerbate atherosclerosis remains unclear.

Expressed on the cell membrane, solute carrier family 7 member 11 (SLC7A11) is an amino acid transporter with its main function being the promotion of cystine uptake and glutathione (GSH) biosynthesis. With GSH functioning as an endogenous antioxidant, homeostatic levels of SLC7A11 protect cells against oxidative stress and ferroptosis (i.e., cell death mediated by iron-dependent lipid peroxidation).^21,22^ As a ferroptosis marker, SLC7A11 plays critical roles in glutamine-GSH metabolism and thus is involved in the pathogenesis of cancer and atherosclerosis.^23^ Ferroptosis inhibition by ferrostatin-1 alleviated atherosclerosis in high-fat diet (HFD)-fed *Apoe*^-/-^ mice with an attendant decrease in iron accumulation and lipid peroxidation and reversed expression of Slc7a11.^24^ Obese rats fed an HFD also showed lower level of Slc7a11, which was associated with increased serum TNF-α level.^25^ Given that EC dysfunction with an exacerbated redox state causes experimental and clinical atherosclerosis, the impaired SLC7A11–GSH axis may be involved in atherogenesis.^26,27^

Notably, *SLC7A11* mRNA is a putative target of miR-30, which prompted us to investigate whether adipocyte-derived miR-30e can target *SLC7A11* in ECs in an exosome-dependent manner.^28^ If true, this endocrine-like mechanism may mediate, in part, obese-aggravated atherosclerosis in experimental animals and humans. Our results show elevated level of miR-30e-5p in adipose-derived exosomes and serum from obese individuals with coronary artery disease (CAD), which was associated with atherosclerosis progression. Moreover, adipose-originated miR-30e-5p could downregulate SLC7A11 and mitochondrial function in ECs. These findings provide new insights into adipocyte–EC crosstalk and their implications in miR-30e antagonism to ameliorate obesity-enhanced atherosclerosis.

## Materials and Methods

### Data availability

The data that supports the findings of this study are available from the corresponding author upon request.

Detailed description of the materials and methods is provided in supplementary material. All clinical samples were obtained at the First Affiliated Hospital of Xi’an Jiaotong University with informed consent and institutional ethics committee approval. Animal experiments were approved by the Institutional Animal Ethics Committee of Xi’an Jiaotong University.

## Results

### Elevated level of adipose-derived miR-30e-5p in obese individuals with CAD

We initially mined published GEO datasets to identify putative miRNAs that link obese and atherosclerosis. Level of miR-30e-5p was significantly increased in serum from atherosclerotic human patients (GSE96621) and circulating miRNA from HFD-fed mice.^29^ However, miR-30e-5p level was lower in serum exosomes from ADicer knockout mice in which miRNA processing is hindered due to Dicer specific ablation in adipose tissue (Figure 1A).^12^ Reanalyzing the GSE25470 dataset, we found that the level of miR-30e-5p was also increased in white adipose tissue (WAT) from obese patients (Figure 1B). To further investigate the causality between obesity, circulating miR-30e-5p, and atherosclerotic disease, we collected serum from CAD patients and healthy controls (HCs) (demographics in Table SII). In agreement with *in silico* analyses of data by others (Figure 1A, 1B), CAD patients registered in this study had higher levels of miR-30e-5p than HCs (Figure 1C). The receiver operating characteristic (ROC) curve revealed that patients would have higher probability of CAD if their miR-30e-5p level was 47.69% higher than the HCs, with sensitivity 78.46% and specificity 69.23% (area under the ROC curve, 0.81) (Figure 1D). We further classified CAD patients into three groups according to their body mass index (BMI): normal weight (BMI ≤ 24.9 kg/m^2^), overweight (25–29.9 kg/m^2^), and obese (≥ 30 kg/m^2^). Serum level of miR-30e-5p was elevated more in obese individuals with CAD (Figure 1E). Furthermore, the circulatory levels of miR-30e-5p were positively correlated with BMI and TG and LDL-C level but negatively with HDL-C level (Figure 1F). Exosomes were then isolated from adipose tissues and serum of obese/CAD individuals as well as non-diseased controls for further analysis (demographics in Table SIII). On nanoparticle tracking analysis, these isolated exosomes had the typical size (≈100 nm) (Figure 1G), and electron microscopy revealed the cup-shaped morphology (Figure 1H). The two groups of exosomes were then labeled with PKH67 and incubated with human aortic endothelial cells (HAECs) *in vitro* (Figure 1I). Level of miR-30e-5p was increased in HAECs co-cultured with exosomes from adipose-tissues or serum of obese/CAD patients (Figure 1J). In line with data from human patients, miR-30e-5p levels were increased in WAT, serum exosomes, and aortic tissues of *ob/ob* mice, *Apoe*^-/-^ mice fed an HFD, and C57BL/6 mice administered AAV8-*PCSK9* and fed an HFD (Figure 1K-M). However, miR-30e-5p level was not elevated in brown adipose tissue (BAT) of these three mouse models (Figure 1K-M). Collectively, results in Figure 1 suggest that the WAT-derived exosomes might transport miR-30e-5p from adipocytes to vascular endothelium in obese humans with CAD, obese mice, and atherosclerotic mice.

**Figure 1.**
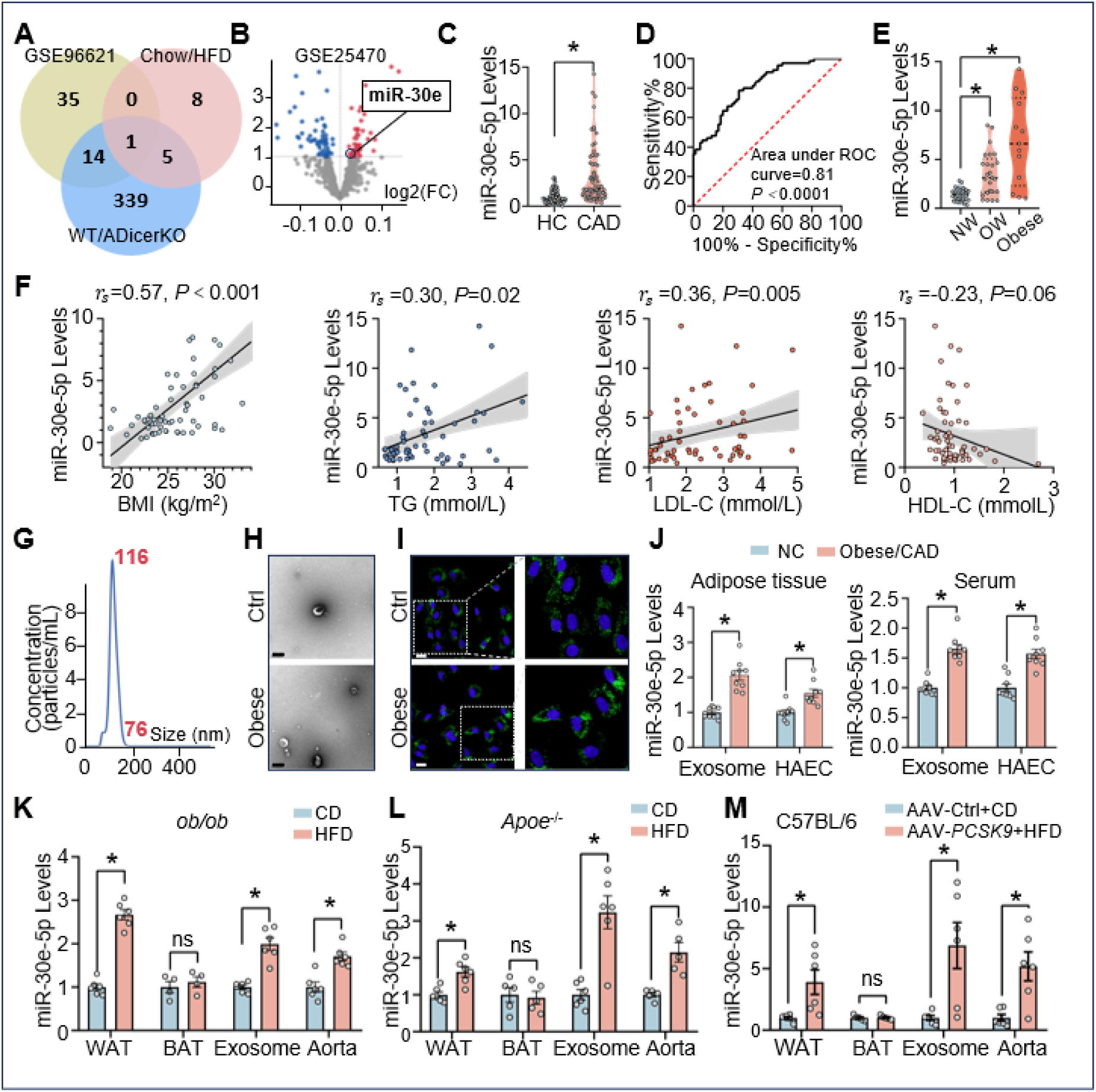
Elevated level of adipose-derived miR-30e-5p in obese individuals with CAD. (**A**) Venn diagram demonstrating the intersection of miRNA levels among 3 databases: miR-30e level was increased in serum from atherosclerotic human patients (GSE96621, chrome yellow) and high-fat diet (HFD)–fed mice (pink) but decreased in serum exosomes from fat-specific Dicer (ADicer)-knockout mice (blue). (**B**) Volcano plot showing increased level of miR-30e in WAT from obese patients (n=30) and normal controls (NCs, n=26) (GSE25470). (**C**) Serum was collected from CAD patients (n=65) and healthy controls (HCs) (n=65) registered in the First Affiliated Hospital of Xi’an Jiaotong University. Serum level of miR-30e-5p was measured by qPCR. The data are fold-change normalized to the mean level of HCs. (**D**) ROC curve with sensitivity and specificity of serum level of miR-30e-5p for differentiating CAD patients from HCs at diagnosis. (**E**) Serum level of miR-30e-5p in CAD patients with normal weight (NW) (BMI ≤ 24.9 kg/m^2^, n=27), overweight (OE) (25–29.9 kg/m^2^, n=25), and obesity (≥ 30 kg/m^2^, n=13). The data are fold-change normalized to the mean of patients with normal weight (NW). (**F**) Correlations between miR-30-5p level and BMI, TG, LDL-C, and HDL-C in CAD patients. (**G**) Size distribution of serum exosomes assessed by nanoparticle tracking analysis. (**H**) Electron microscopy images of adipose tissue-derived exosomes, approximately 100 nm in diameter. Scale bar: 100 nm. (**I**) Serum exosomes were labeled with PKH67 green fluorescence dye, incubated with HAECs for 24 h, then fixed for confocal imaging. Nuclei were counterstained with DAPI (blue). Scale bar: 20 μm. (**J**) HAECs were incubated with exosomes isolated from subcutaneous adipose tissues or serum from obese patients with CAD or non-diseased controls for 24 hr. qPCR analysis of miR-30e-5p level in exosomes and co-cultured HAECs. (**K-M**) Levels of miR-30e-5p in WAT, brown adipose tissue (BAT), serum exosomes, and aortic intima from *ob/ob* mice fed a chow diet or HFD for 8 weeks (K); *Apoe*^-/-^ mice fed with chow diet or HFD for 12 weeks (L); and C57BL/6 mice injected with AAV8-Ctrl virus via the tail vein and fed a chow diet (AAV8-Ctrl+CD) or administered AAV8-*PCSK9* virus and fed an HFD (AAV8-*PCSK9*+HFD) for 12 weeks (M). Data are mean±SEM from 4-6 mice per group. Normally distributed data were analyzed by 2-tailed Student *t*-test (aorta in K) or 2-tailed Student *t*-test with Welch correction [WAT in K; WAT, exosome, and aorta in L; WAT and exosomes in (M)] and non-normally distributed data by Mann-Whitney U test [(C) and (J); BAT and exosome in (K); BAT in (L); BAT and aorta in (M)] between the 2 indicated groups. Data were analyzed by Brown-Forsythe ANOVA test with Dunnett’s multiple comparisons test among multiple groups in (E). Spearman analysis revealing correlations between miR-30-5p and BMI and TG, LDL-C, and HDL-C levels in (F). \**P* < 0.05. ns=not significant.

### MiR-30e-5p targets SLC7A11

Having established that adipose tissue-derived miR-30e-5p might be an endocrine-like miRNA regulating adipose-vessel crosstalk, we then investigated the downstream target of miR-30e-5p. For an unbiased screening, we performed RNA-seq analysis of HAECs transfected with miR-30e-5p or control miRNA. Gene Ontology (GO) analysis of differentially regulated pathways showed that miR-30e-5p level was correlated nitric oxide biosynthetic process, glutathione metabolic process, regulation of fat cell differentiation, cellular oxidant detoxification, lipids and atherosclerosis, and SLC-mediated transmembrane transport (Figure 2A). The heatmap shown in Figure 2B reveals that miR-30e-5p overexpression downregulated genes involved in oxidant detoxification, glutathione metabolic process, and nitric oxide biosynthesis. In addition, miR-30e-5p overexpression affected genes involved in lipid and atherosclerosis as well as SLC-mediated transmembrane transport. Markedly, miR-30e-5p downregulated atheroprotective genes (e.g., SOD2) and upregulated atheroprone genes (e.g., NOX4) involved in lipid and atherosclerosis. For SLC family transporters, miR-30e-5p downregulated SLC7A11 and upregulated SLC2A3 (GLUT3), which have been shown to affect cellular redox state.^30^ Using TargetScan, we predicted that miR-30e-5p could target 3′-UTR of *SLC7A11* mRNA (Supplemental Figure 1A). SLC7A11 level was reduced in dysfunctional ECs due to lipid accumulation (GSE246083) and in ECs exposed to atheroprone flow (GSE118717).^31,32^ These *in silico* analyses showing miR-30e-5p targeting *SLC7A11* were depicted by Venn analysis in Figure 2C. According to functional roles of SLC7A11 in cell death and the glutathione redox system (Figure 2D), we theorized that miR-30e-5p targets the 3′-UTR of *SLC7A11* mRNA (Supplemental Figure 1A). Of note, miR-30e-5p and its targeted sequences in the 3′-UTR of *SLC7A11* mRNA are highly conserved in various mammalian species, including human, mouse, rat, chimp, rhesus, pig, and cow (Supplemental Figure 1A). We then constructed luciferase reporters fused with the *SLC7A11*-3′-UTR (WT) or mutated *SLC7A11*-3′-UTR (MT) to test whether miR-30e-5p directly targets the *SLC7A11* mRNA (Figure 2E). Overexpression of miR-30e-5p suppressed the *SLC7A11*-3′-UTR (WT) luciferase activity but not MT activity (Figure 2E). Furthermore, HAECs transfected with miR-30e-5p had lower protein and mRNA levels of SLC7A11 (Figure 2F, Supplemental Figure 1B). In contrast, inhibition of miR-30e-5p increased levels of SLC7A11 and its mRNA, in a *yin-yang* manner (Figure 2G, Supplemental Figure 1C). Furthermore, miR-30e-5p and *SLC7A11* mRNA levels were enriched in Ago1 and Ago2, namely, the miRNA-induced silencing complexes (miRISCs), in HAECs transfected with miR-30e-5p (Figure 2H), so *SLC7A11* mRNA was a bona fide target of miR-30e-5p in ECs. Because SLC7A11 regulates the exchange of glutamate and cystine, we analyzed metabolites in HAECs co-cultured with adipocytes that were stimulated with palmitate and transfected with or without miR-30e-5p inhibitor. As anticipated, the co-cultured HAECs and the secreted exosomes had an increased level of miR-30e-5p, and miR-30e-5p inhibitor abolished this increase (Supplemental Figure 1D). The untargeted metabolomics depicted by KEGG showed that glutamate and glutathione metabolism were significantly enriched (Figure 2I). In line with these metabolomic changes, HAECs co-cultured with exosomes isolated from palmitate-stimulated adipocytes showed an increased level of glutamate and a decreased level of cystine. These glutamate/cystine alterations were reversed by treatment with the miR-30e-5p inhibitor (Figure 2J). Importantly, incubation of HAECs with serum from obese patients with CAD also led to increased glutamate and decreased cystine levels (Figure 2K). In line with these results, HAECs treated with serum from obese CAD patients showed reduced cystine uptake, which was rectified by miR-30e-5p inhibition (Figure 2L). Furthermore, given that miR-30e-5p targets *SLC7A11* mRNA, adenovirus-mediated SLC7A11 overexpression rescued the cystine uptake by ECs (Figure 2M).

**Figure 2.**
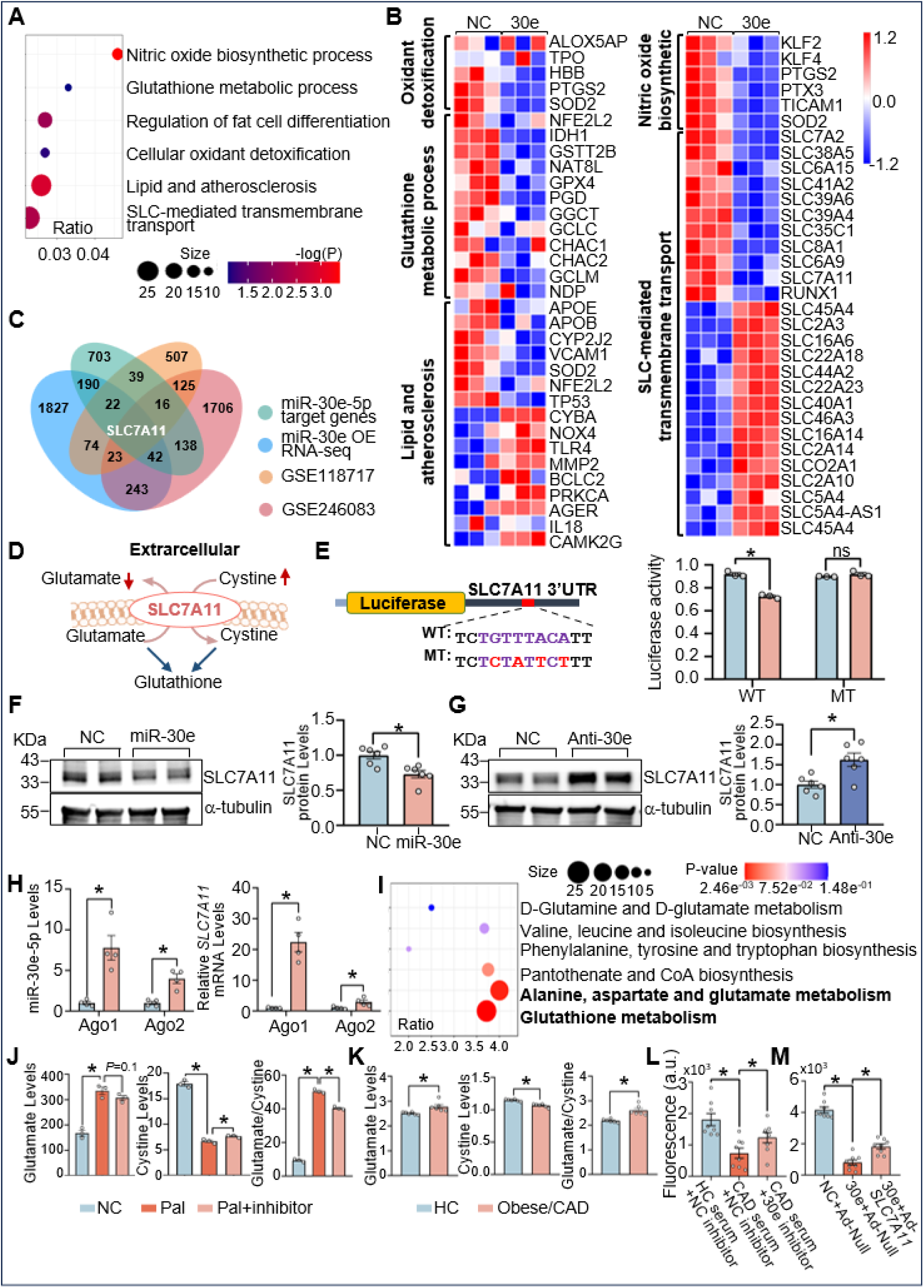
MiR-30e-5p targets SLC7A11. (A, B) HAECs were transfected with miR-30e-5p or control miRNA (NC) for 48 h. RNAs were isolated and subjected to RNA-seq analysis. GO enrichment was delineated by Metascape for the 1690 differentially expressed genes with the cutoff of log2 fold-change > 1 or <-1, padj < 0.1, and plotted according to gene ratio, size and-log (*P*-value) of every term (A). Heatmap comparison of log2 fold-change of the indicated differentially expressed genes according to *Z*-scores (B). (**C**) Venn diagram showing suppressed SLC7A11 as a target of miR-30e-5p (https://www.targetscan.org/vert_80/, light green) in lipid-accumulated ECs (GSE246083, pink), ECs exposed to atheroprone flow (GSE118717, orange), and miR-30e-5p overexpression (miR-30e OE RNA-seq, blue). (**D**) Graphic presentation of the SLC7A11-mediated glutamate/cystine metabolism. (**E**) HAECs were co-transfected with miR-30e-5p and a dual luciferase reporter fused with the 3’-UTR of *SLC7A11* (Luc-*SLC7A11*-3’-UTR-WT) or a mutant reporter with miR-30e-5p binding site mutated (Luc-*SLC7A11*-3’-UTR MT). The firefly and Renilla luciferase activity were measured (n=3). (**F, G**) HAECs were transfected with miR-30e-5p or control miRNA (F), and miR-30e-5p inhibitor (Anti-30e) or control miRNA (G). SLC7A11 was measured by western blot analysis (n=6). (**H**) HAECs were transfected with miR-30e-5p or NCs. Ago-1 or Ago-2 were immunoprecipitated from the transfected ECs. The miRISC-associated miR-30e-5p and *SLC7A11* mRNA were quantified by qPCR (n=4). (**I, J**) 3T3L1-derived adipocytes were incubated with or without palmitate (400 mM) for 24 h (Pal; NC). The palmitate-incubated adipocytes were also transfected with miR-30e-5p inhibitor (Pal+inhibitor). n=3 in each of the three cell groups. HAECs were incubated with exosomes isolated from the three groups of adipocytes, followed by untargeted metabolomic analysis. (**I**) KEGG enrichment pathways of metabolic variations among NCs, Pal, and Pal+inhibitor groups, *P*<0.05. (**J**) Changes in glutamate, cystine, and the glutamate/cystine ratio among these three groups. (**K**) HAECs were incubated with serum from obese patients with CAD or healthy controls (HC). Levels of glutamate and cystine in ECs were measured by ELISA. (**L**) HAECs were transfected with miR-30e-5p inhibitor or control miRNA (NC). Transfected cells were then treated with serum from obese patients with CAD or healthy controls. (**M**) HAECs were transfected with miR-30e-5p or NCs, then infected with Ad-*SLC7A11* or Ad-Null (10 MOI). Cystine uptake by HAECs in (L, M) was assessed accordingly (n=8). Normally distributed data were analyzed by 2-tailed Student *t*-test (E, F, G, cystine in K) or Welch’s *t*-test (glutamate and the glutamate/cystine ratio in K) for the indicated groups. Non-normally distributed data in (H) were analyzed by Mann-Whitney *U* test. Data in (J, L, M) were analyzed by 2-way ANOVA with Holm-Šídák post *hoc* test among multiple groups. \**P* < 0.05.

### SLC7A11 is decreased in experimental atherosclerotic mice and CAD patients

To explore the crosstalk between adipocytes and vascular endothelium in the context of miR-30e-5p–SLC7A11 and atherogenesis, we performed single-nucleus RNA-sequencing (snRNA-seq) of WAT from *Apoe*^-/-^ mice fed an HFD or chow diet. On tSNE analysis, we identified 5 cell clusters, including adipocytes, epithelial cells, fibroblasts, ECs, and immune cells (Figure 3A), according to their distinct gene expression profiles (Supplemental Figure 2A). HFD increased the number of pro-inflammatory macrophages, as signified by increased level of *Tnf*, *Nlrp3*, *Cd74*, etc. (Supplemental Figure 2B). In adipocytes and ECs, HFD reduced the *Slc7a11* mRNA level (Figure 3B, 3C). Pseudotime trajectory analysis demonstrated that *Slc7a11* expression was higher in early stages of adipocyte and EC subpopulations (state 1) and decreased afterwards to the lowest level in state 5. Moreover, atherosclerotic tissues had the highest level of state 5 cells (Figure 3D, 3E). These results suggest SLC7A11 suppression during the atherogenic course and an inverse association between SLC7A11 abundance and EC homeostasis. In line with this notion, mRNA and protein levels of Slc7a11 were reduced in the aortic intima of atherosclerotic *Apoe*^-/-^ mice (Figure 3F). Slc7a11 level reduction in aortic intima was recapitulated in C57BL/6 mice administered AAV8-*PCSK9* and fed an HFD (Figure 3G). For human atherosclerosis, we performed immunostaining experiments to examine the level of SLC7A11 in atherosclerotic plaque from patients who underwent carotid endarterectomy (demographics Table SIV). Shown in Figure 3H, SLC7A11 expression was reduced in atherosclerotic lesions as compared with non–diseased specimens. To explore changes in metabolites resulting from the differential expression of SLC7A11, we performed untargeted metabolomics analysis of serum from CAD patients and non-diseased controls (NCs) in a discovery cohort (demographics Table SV). KEGG enrichment analysis revealed that glutamate metabolism, cysteine and methionine metabolism, and glutathione metabolism varied the most between CAD patients and NCs (Figure 3I). With SLC7A11 mediating the extracellular influx of cysteine in the expense of efflux of intracellular glutamate, we studied the serum metabolites in a validation cohort (demographics Table SVI). The lower level of glutamate associated with a higher level of cystine was confirmed in CAD patients in the discovery and validation cohorts (Figure 3J, 3L). However, the serum level of glutamate was negatively correlated with BMI (Figure 3K, 3M). Moreover, ROC curves in Figure 3N showed that patients with glutamate levels < 37.93% and 31.82% of those in NCs might have CAD, with sensitivity 41.38% and 40.91% and specificity 96.55% and 90.91% in the discovery and validation cohorts, respectively (area under the ROC curve: 0.68 in the discovery cohort; 0.69 in the validation cohort). Taken together, data in Figure 3 suggest that SLC7A11 was downregulated in the vascular endothelium in patients with CAD as well as mice with experimental atherosclerosis, thus increasing glutamate level but decreasing cystine level in the circulation.

**Figure 3.**
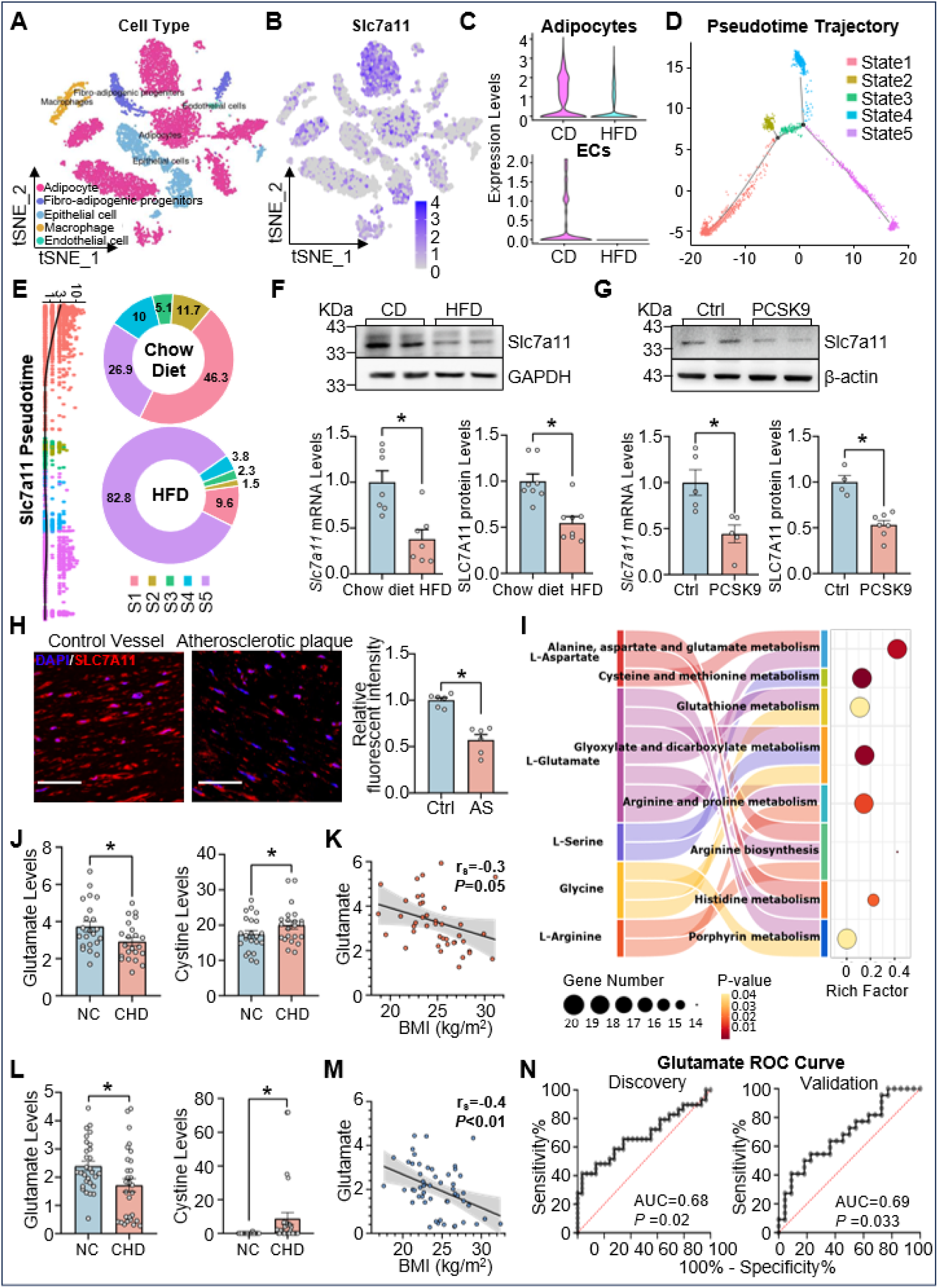
SLC7A11 is decreased in experimental atherosclerotic mice and CAD patients. (**A-E**) Eight-week-old male *Apoe*^-/-^ mice were fed an HFD or chow diet for 12 weeks. snRNA-seq was conducted on epididymal WAT (eWAT) collected from 3 animals in each of the two groups, which generated transcriptomics from 9540 heterogenous cells. (**A**) tSNE clustering showing 5 annotated cell types in eWAT. (**B**) Biaxial scatter plot displaying levels of *Slc7a11* in these 5 cell types, with grey and purple representing low and high levels of *Slc7a11*. (**C**) Violin plots showing levels of *Slc7a11* in annotated adipocytes and ECs. (**D**) Cell development pseudotime trajectory plot of nuclei from eWAT, showing 5 evolving stages. (**E**) The percentage of cells at each of the 5 stages and the vertical plot on the left reveals the dynamics of *Slc7a11* expression at different stages with each point as a single nucleus. (**F**) Slc7a11 protein and mRNA levels in the aortic intima in *Apoe*^-/-^ mice fed an HFD or chow diet. n=7 for qPCR; n=8 for western blot analysis. (**G**) Eight-week-old C57BL/6 mice were administered a single dose of AAV8-*PCSK9* or AAV8-Ctrl (1×10^11^ vg/mouse) by tail vein injection and fed an HFD or chow diet for 12 weeks. Aortic intima protein and mRNA levels of Slc7a11 were assessed. n=5 for qPCR; n=4-7 for western blot analysis. (**H**) Representative immunofluorescence staining of SLC7A11 in human carotid atherosclerotic plaques obtained from carotid endarterectomy and control abdominal vessels from autopsy. Specimens were counterstained with DAPI (blue). n=3, Scale bar: 50 μm. (**I, J**) Untargeted metabolomics analysis of serum from CAD patients and NCs in the discovery cohort. (I) KEGG enrichment pathway analysis based on differential metabolites between 2 groups (fold change >1.5 or <-1.5, P<0.05). (J) Variations in glutamate and cystine levels between CAD patients and NCs. (**K**) Correlation between serum glutamate level and BMI. (**L, M**) Targeted metabolomics analysis of serum glutamate and cystine levels in validation cohort, confirming the inverse correlation between serum glutamate level and BMI. (**N**) ROC curve with sensitivity and specificity of serum level of glutamate in discovery and validation cohorts, differentiating CAD patients from NCs. Data were analyzed by 2-tailed Student *t*-test (F, qPCR in G, H); Mann-Whitney *U* test between 2 indicated groups (WB in G, glutamate and cystine in J, L); Spearman analysis in (K) and Pearson analysis in (M) demonstrate negative correlations between serum glutamate levels and BMI. \**P* < 0.05.

### MiR-30e-5p–SLC7A11 regulates central carbon metabolism and mitochondrial function

With the established role of SLC7A11 in cystine uptake and biosynthesis, we then explored the effect of SLC7A11 on EC metabolomic changes by using LC-MS/MS analysis to compare metabolites in HAECs infected with *SLC7A11* adenovirus (Ad-*SLC7A11*) versus Ad-Null. SLC7A11 overexpression significantly increased levels of 29 metabolites in ECs versus Ad-Null–infected control cells (Figure 4A, Table SVII). Pathway enrichment analysis revealed that the pathways including central carbon metabolism in cancer, carbon metabolism, ABC transporters, biosynthesis of amino acids, aminoacyl-tRNA biosynthesis and alanine, aspartate and glutamate metabolism were significantly changed by SLC7A11 overexpression (Figure 4B). Because glycolysis and the tricarboxylic acid (TCA) cycle are two crucial pathways in central carbon metabolism, we used Seahorse assays to investigate the role of SLC7A11 in regulating EC glycolysis and oxidative phosphorylation. SLC7A11 overexpression in ECs decreased ECAR but increased OCR (Figure 4C, 4D). By contrast, SLC7A11 knockdown increased ECAR and decreased OCR (Supplemental Figure 3A, 3B). Consistent with miR-30e-5p targeting *SLC7A11* mRNA, ASO against miR-30e-5p increased OCR and decreased ECAR in ECs (Supplemental Figure 3C, 3D). Furthermore, SLC7A11 overexpression reversed the miR-30e-5p–increased ECAR and –decreased OCR (Figure 4E, 4F). These results suggest that the miR-30e-5p–SLC7A11 axis impaired carbon metabolism in ECs. Because of the negative effect of this axis on oxidative phosphorylation, we studied whether it regulates EC mitochondrial function. On confocal imaging using MitoTracker, HAECs transfected with miR-30e-5p showed decreased mitochondria in ECs and increased mitochondrial fragmentation, which was rectified by SLC7A11 overexpression (Figure 4G). These results led us to theorize that this endocrine-like axis causes EC dysfunction. In agreement, SLC7A11 knockdown in ECs increased the expression of *FN1* and *SNAI1* but decreased that of *CDH5* and *NR2F2* (Figure 4H), indicative of augmented endothelial-to-mesenchymal transition (EndoMT). Regarding redox stress, SLC7A11 knockdown increased ROS levels in HAECs, and SLC7A11 overexpression reversed the miR-30e–increased ROS production (Figure 4I, 4J). In line with these EC dysfunctional events, SLC7A11 deficiency decreased eNOS Ser-1177 phosphorylation, thereby manifesting the reduced eNOS-derived NO bioavailability (Supplemental Figure 3E).

**Figure 4.**
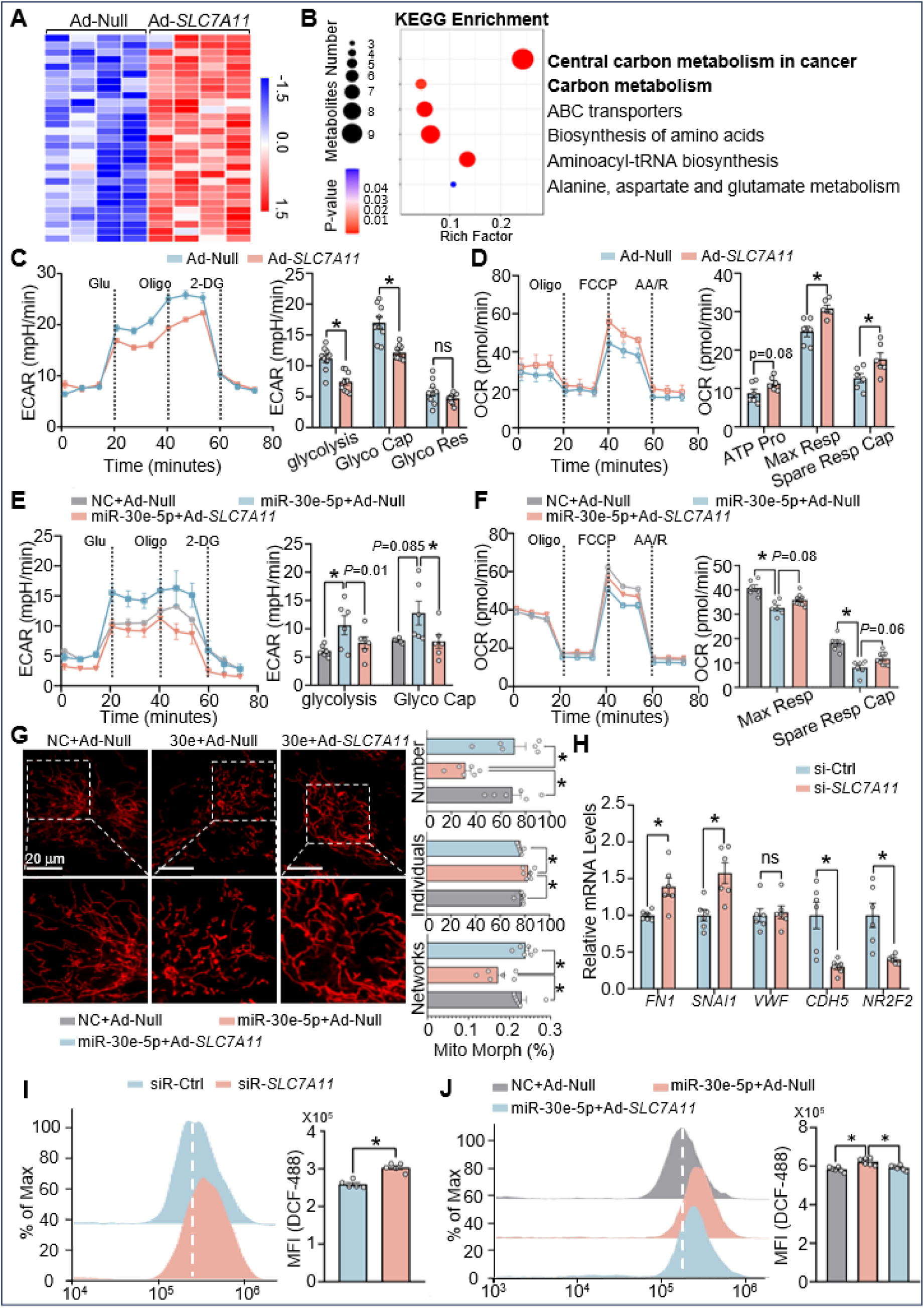
MiR-30e-5p–SLC7A11 regulates central carbon metabolism and mitochondrial function. (**A, B**) HAECs were infected with Ad-*SLC7A11* or Ad-Null for 48 hr, then analyzed by untargeted metabolomics (n=4). Fold change ≥1.2 or ≤ 0.83 and *P* < 0.05 were used as the screening criteria. (**A**) Heatmap showing metabolites differentially regulated by SLC7A11 overexpression. (**B**) KEGG enrichment analysis of the 29 upregulated metabolites. (**C-F**) Extracellular acidification rate (ECAR) and oxygen consumption rate (OCR) assays of HAECs infected with Ad-*SLC7A11* (10 MOI) or Ad-Null (C,D; n=9); and transfected with or without NC miRNA (NC), miR-30-5p and with or without Ad-*SLC7A11*, as indicated (E,F; n=6-9). (**G**) Representative confocal imaging with MitoTracker for mitochondrial morphology in HAECs transfected with NC miRNA or miR-30e-5p and infected with Ad-*SLC7A11* or Ad-Null (n=6, 10 cells counted for each replicate). Scale bar=10 μm. (**H,I**) HAECs were transfected with siR-*SLC7A11* or siR-Ctrl. Levels of indicated mRNAs were determined (n=6) in (H); FACS detection of oxidized DCF (n=5) in (I). (**J**) HAECs were transfected with NC miRNA or miR-30e-5p and infected with Ad-*SLC7A11* or Ad-Null, then analyzed by FACS detection of oxidized DCF (n=6). Normally distributed data (glycolysis in C; D; *SNAI1* and *VWF* in H; I) were analyzed by 2-tailed Student *t*-test, or (Gylco Cap and Gylco Res in C; *FN1*, *CDH5* and *NR2F2* in H) by Welch’s *t*-test. Data among multiple groups in (E, F, G, J) were analyzed by two-way ANOVA with the Holm-Šídák post *hoc* test. \**P* < 0.05. ns=not significant. Glu, glucose; Oligo, Oligomycin; 2-DG, 2-dexoy-D-lucose; FCCP, carbonyl cyanide-p-(triffuoromethoxy)phenylhydrazone; AA/R, antimycin A& rotenone; Glyco Cap, glycolytic capacity; Glyco Res, glycolytic reserve; ATP Pro, ATP production; Max Resp, maximal respiration; Spare Resp Cap, spare respiration capacity.

### EC-specific *Slc7a11* deficiency aggravates atherosclerosis in mice

To determine the atherogenic effect resulting from the miR-30e–SLC7A11 axis, we created EC-*Slc7a11*^-/-^ mice. Endothelial ablation of *Slc7a11* in this line of mice was validated in isolated pulmonary ECs (Figure 5A). To induce atherosclerosis, 8-week-old male and female EC-*Slc7a11*^-/-^ mice (ECKO) and their *Slc7a11*^flox/flox^ wild type (WT) littermates were administered AAV8-*PCSK9* via tail vein injection, then fed an HFD for 12 weeks before being euthanized (Figure 5B). Although lipid profiles, including TC, TG, LDL-C, and HDL-C levels, did not significantly differ among EC-*Slc7a11*^-/-^ and *Slc7a11*^flox/flox^ littermates (Figure 5C), *en face* Oil-red O staining revealed a significant increase in atherosclerotic lesions in the aortas of EC-*Slc7a11*^-/-^ male and female mice as compared with *Slc7a11*^flox/flox^ littermates (Figure 5D). However, control EC-*Slc7a11*^-/-^ or *Slc7a11*^flox/flox^ mice fed a chow diet for 12 weeks showed little, if any, atherosclerosis (Figure 5D). Increased Oil-red O and Cd68 staining were also evident in the aortic roots of EC-*Slc7a11*^-/-^ mice, thereby reinforcing that *Slc7a11* ablation enhanced the atherosclerosis susceptibility (Figure 5E). Of note, EC-*Slc7a11*^-/-^ mice showed decreased serum level of glutamate and increased level of cystine as compared with *Slc7a11*^flox/flox^ littermates (Figure 5F, 5G), which suggests detrimental effects because of *Slc7a11* deficiency in the vascular endothelium.

**Figure 5.**
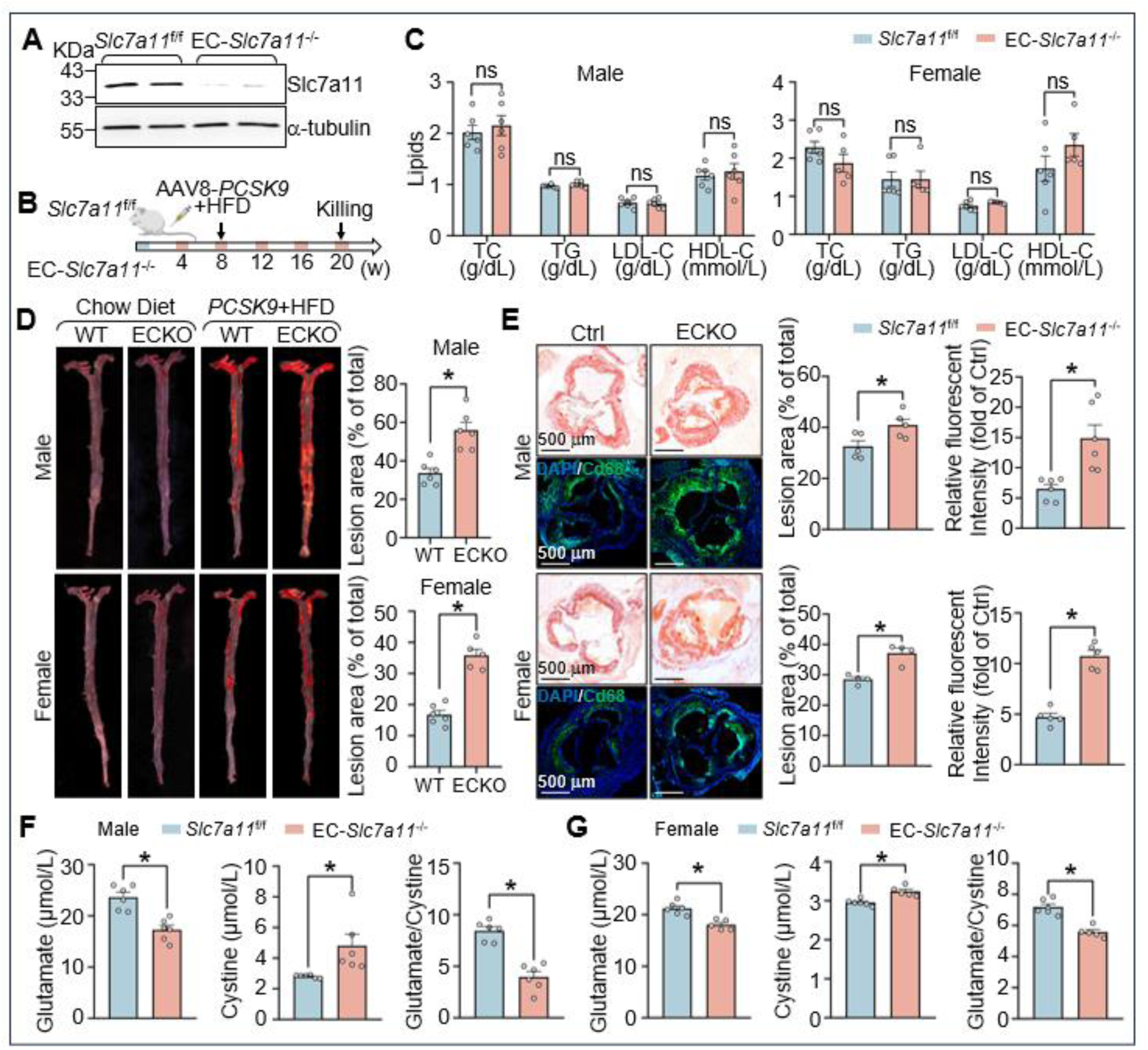
**EC-specific *Slc7a11* deficiency aggravates atherosclerosis in mice**. (**A**) Western blot analysis showing Slc7a11 and α-tubulin in lung ECs from EC-*Slc7a11*^-/-^ and their *Slc7a11*^flox/flox^ control littermates. (**B**) Eight-week-old male and female EC-*Slc7a11*^-/-^ and their *Slc7a11*^flox/flox^ littermates were administered a single dose of AAV8-*PCSK9* (1×10^11^ vg/mouse) via tail vein injection and fed an HFD for 12 weeks. (**C**) Levels of serum TC, TG, LDL-C, and HDL-C in the two groups of mice. (**D**) *En face* Oil-red O staining of aortic specimens. (**E**) Histological sections of aortic roots stained with Oil-red O (atherosclerosis) or immunostaining with anti-Cd68 antibody (macrophage, green). The specimens were counterstained with DAPI (blue). Scale bars: 500 μm. (**F,G**) Levels of serum glutamate and cystine and their ratios in the two groups of mice. Data are mean±SEM from 4-6 mice per group. Normally distributed data were analyzed by 2-tailed Student *t*-test (male in C and D; glutamate and glutamate/cystine ratio in F) or Welch’s *t*-test (male Cd68 in E; cystine in F). Non-normally distributed data (female in C, D and E; male Oil-red O in E; G) were analyzed by Mann-Whitney *U* test. \**P* < 0.05. ns=not significant.

### Exogenous miR-30e-5p exacerbates atherosclerosis

With the atheroprone role of SLC7A11 deficiency deduced in Figure 5, we then tested whether exogenously delivered miR-30e-5p increases atherosclerosis, like that with SLC7A11 deficiency. For this, we administered *Apoe*^-/-^ mice miR-30e-5p or control miRNA via tail-vein injection, then fed these mice an HFD for 12 weeks (Figure 6A, Supplemental Figure 4A). The level of miR-30e-5p was significantly elevated in serum, aortas, and WAT of *Apoe*^-/-^ mice receiving miR-30e-5p as compared with mice receiving control miRNA (Figure 6B, 6C, Supplemental Figure 4B). The miR-30e-5p experimental and control mice did not differ in body weight and TC, TG, LDL-C, and HDL-C levels (Figure 6D, 6E). Corresponding to the miR-30e-5p administration in the experimental group, the aortic level of *Slc7a11* mRNA was reduced and the circulatory level of glutamate decreased and level of cystine increased (Figure 6F, 6G). At the disease level, miR-30e-5p administration significantly increased atherosclerosis compared to mice receiving control miRNA (Figure 6H, 6I, Supplemental Figure 4C-E). Together, the results in Figure 6 show that the exogenously delivered miR-30e-5p, mimicking the obesity-generated miR-30e-5p, aggravated atherosclerosis in *Apoe*^-/-^ mice.

**Figure 6.**
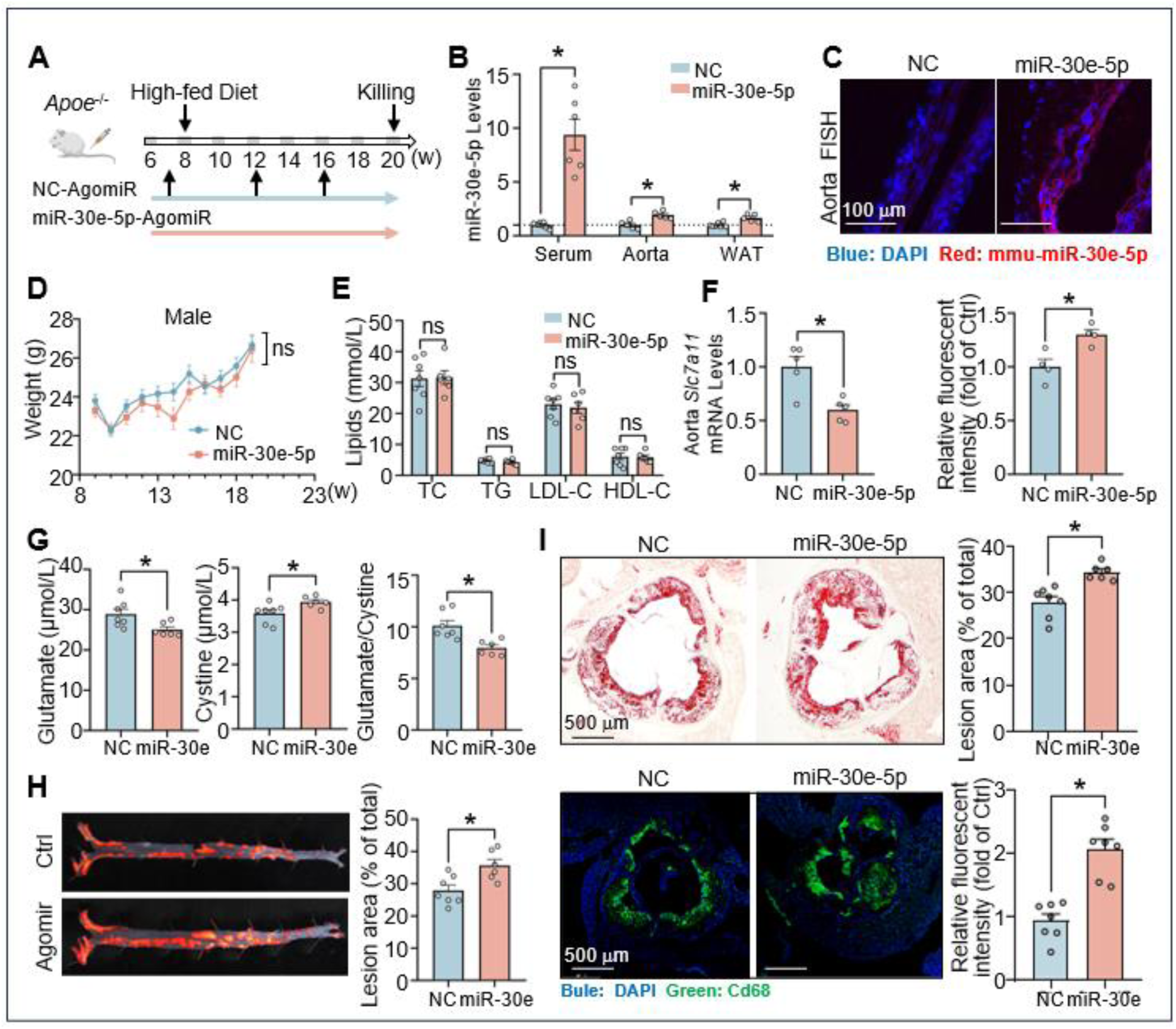
Exogenous miR-30e-5p exacerbates atherosclerosis. (**A**) At 7 weeks old, male *Apoe*^-/-^ mice received miR-30e-5p-agomir (miR-30e) or NC miRNA via tail-vein injection at 16 mg/kg body weight. At week 8, mice were fed an HFD and the 2^nd^ and 3^rd^ doses of agomir were given 4 and 8 weeks post-HFD. (**B**) qPCR analysis of miR-30e-5p in serum, aortic intima, and WAT from 2 groups of mice. (**C**) FISH assay and confocal microscopic images showing the aortic amount of miR-30e-5p. (**D**) Body weights; (**E**) levels of TC, TG, LDL-C, and HDL-C; (**F**) aortic intima levels of *Slc7a11* mRNA; (**G**) serum levels of glutamate and cystine and their ratios; (**H**) *en face* Oil-red O staining of atherosclerotic lesions. (**I**) Oil-red O and DAPI staining and anti-Cd68 immunostaining of the aortic root of the 2 groups of mice. Data are mean±SEM from 4-7 mice per group. Normally distributed data were analyzed by 2-tailed Student *t*-test (aorta in B; E; F; G; H; I) or Welch’s *t*-test (serum in B) and non-normally distributed data by Mann-Whitney *U* test (C; WAT in B). \**P* < 0.05.

### Atherosclerosis reduction by miR-30e-5p antagonism

Complementary to experiments in Figure 6, we used a miR-30e-5p loss-of-function approach to test whether the exogenously delivered anti-miR-30e-5p could mitigate atherosclerosis in *Apoe*^-/-^ mice (Figure 7A). Accordingly, administration of miR-30e-5p antagomir reduced the level of miR-30e-5p in circulation, aorta, and WAT but increased the *Slc7a11* mRNA level in aortas as compared with *Apoe*^-/-^ mice receiving control miRNA (Figure 7B, 7C, Supplemental Figure 5A). Although body weight and lipid profiles of the two groups of mice were comparable (Figure 7D, 7E, Supplemental Figure 5B), mice receiving miR-30e-5p antagomir showed reduced atherosclerosis (Figure 7F, Supplemental Figure 5C). Given that adipocytes are the putative tissue source of miR-30e-5p, we injected LNP-anti-miR-30e-5p to WAT located in the inguinal area of *ob/ob* mice (Figure 7G). This administration lowered the level of miR-30e-5p in the injected adipose tissues and also in circulation and aortas (Figure 7H-J). It significantly increased the expression of *Slc7a11* in the aortic intima, with attendant reduction of atherosclerosis (Figure 7K). Because *ob/ob* mice administered LNP-anti-miR-30e-5p or LNP-Control did not significantly differ in lipid profile or body weight (Supplemental Figure 5D, 5E), the mechanism by which LNP-anti-miR-30e-5p reduced atherosclerosis would be independent of hyperlipidemia (Figure 7L-M). Rather, LNP-anti-miR-30e-5p administration decreased the circulatory level of glutamate and increased that of cystine as compared with LNP-Ctrl administration (Supplemental Figure 5F). Altogether, results from animal models with EC ablation of SLC7A11 (Figure 5), miR-30e-5p gain-of-function (Figure 6), and miR-30e-5p loss-of-function (Figure 7) indicate that the endocrine-like miR-30e-5p–SLC7A11 axis would contribute to obesity-aggravated atherosclerosis in rodents.

**Figure 7.**
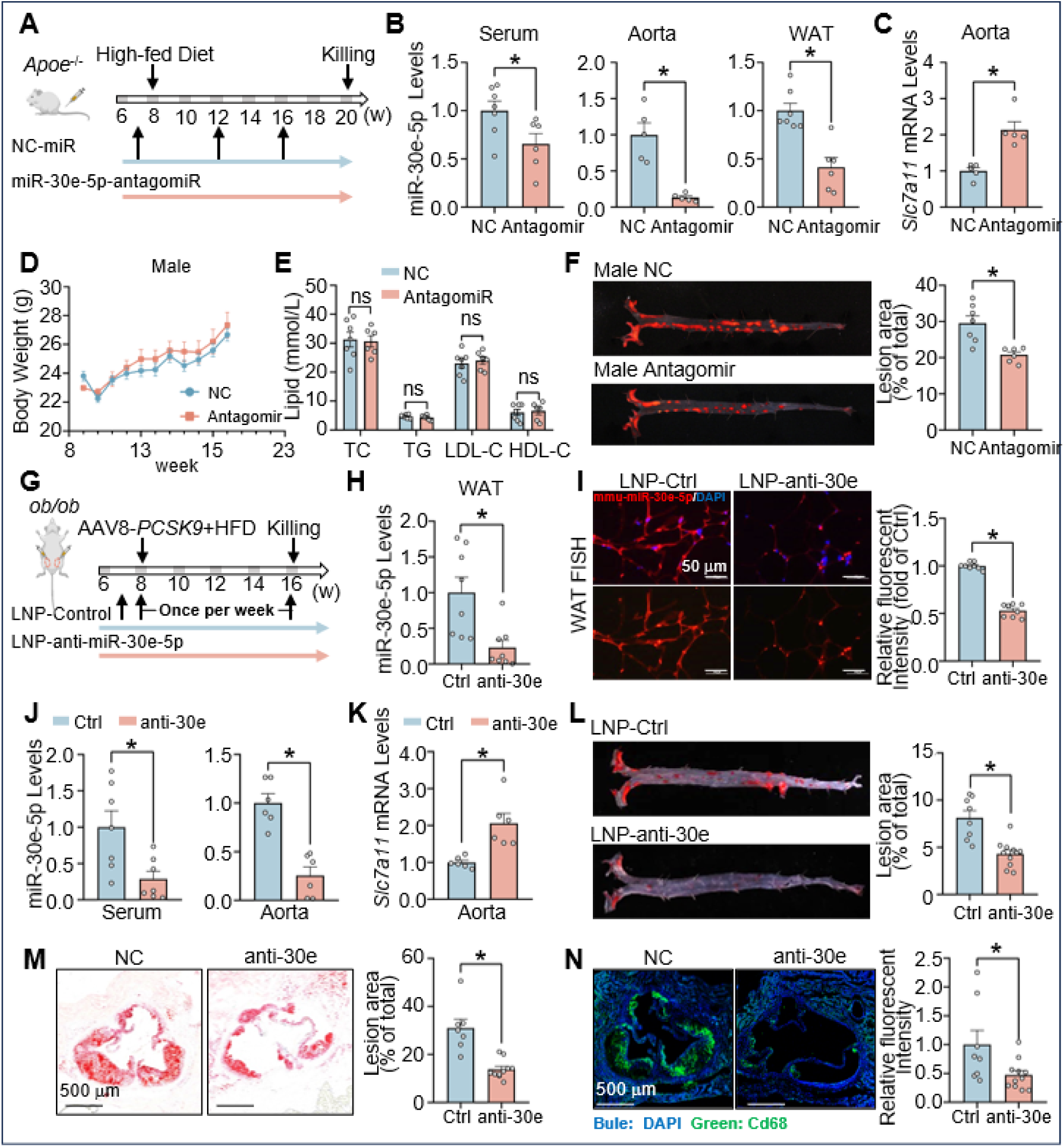
Atherosclerosis reduction by miR-30e-5p antagonism. (**A**) The *Apoe*^-/-^ mouse groups, feeding protocol, and antiagomir delivery were the same as described in Figure 6A. n=7 in NCs and n=6 in miR-30e-5p-antagomir group. (**B-F**) Levels of miR-30e-5p in serum, aortic intima, and WAT (B). Level of *Slc7a11* mRNA in mouse aortic intima (C); body weights (D); levels of serum TC, TG, LDL-C, and HDL-C (E); Oil-red O staining of atherosclerosis (F). (**G**) At 7 weeks old, male *ob/ob* mice received 2 mg/kg/wk LNP-anti-miR-30e-5p or LNP-control miRNA (LNP-Ctrl) in WAT in the inguinal area. One week later, all mice were administered AAV8-*PCSK9* and fed an HFD to induce hyperlipidemia. (**H-N**) Levels of miR-30e-5p in WAT by qPCR (H) and FISH (I) and in serum and aortic intima (J); level of *Slc7a11* mRNA in mouse aortic intima (K); Oil-red O staining of atherosclerosis in aortic specimens and aortic root (L,M); anti-Cd68 immunostaining (N). n=8 in LNP-Ctrl group and n=11 in LNP-anti-miR-30e-5p group. Data are mean±SEM from 6-11 mice per group. Normally distributed data were analyzed by 2-tailed Student *t*-test (I, J; L, M) or Welch’s *t*-test (K) and non-normally distributed data (H, N) by Mann-Whitney *U* test. \**P* < 0.05.

## Discussion

This study elucidates a pathophysiological causality between obesity and atherosclerosis via an endocrine-like mechanism involving the adipocyte-derived miR-30e-5p and its target SLC7A11 in the vascular endothelium. The major findings are (1) elevated circulating level of miR-30e-5p in obese individuals with CAD; (2) decreased SLC7A11 level in CAD patients and experimental atherosclerotic mice; (3) miR-30e-5p directly targeted the 3′-UTR of *SLC7A11* mRNA, resulting in dysfunctional ECs, including dysregulated thioreduction, increased glycolysis, reduced oxidative phosphorylation, and impaired mitochondrial function; and (4) ablation of *Slc7a11* in endothelium aggravated but inhibition of miR-30e-5p in adipose tissue alleviated atherosclerosis in mouse models (illustrated in Figure 8). From these findings, we propose that miR-30e-5p targeting SLC7A11 links the maladapted adipose tissue with atherosclerosis in the arterial wall. Thus, the miR-30e-5p– SLC7A11 axis might be integral to the CKM syndrome. This notion supports the emerging view that adipose tissue, traditionally considered an energy storage depot, is an active endocrine organ that affects vascular health via secreted factors.^33^

**Figure 8.**
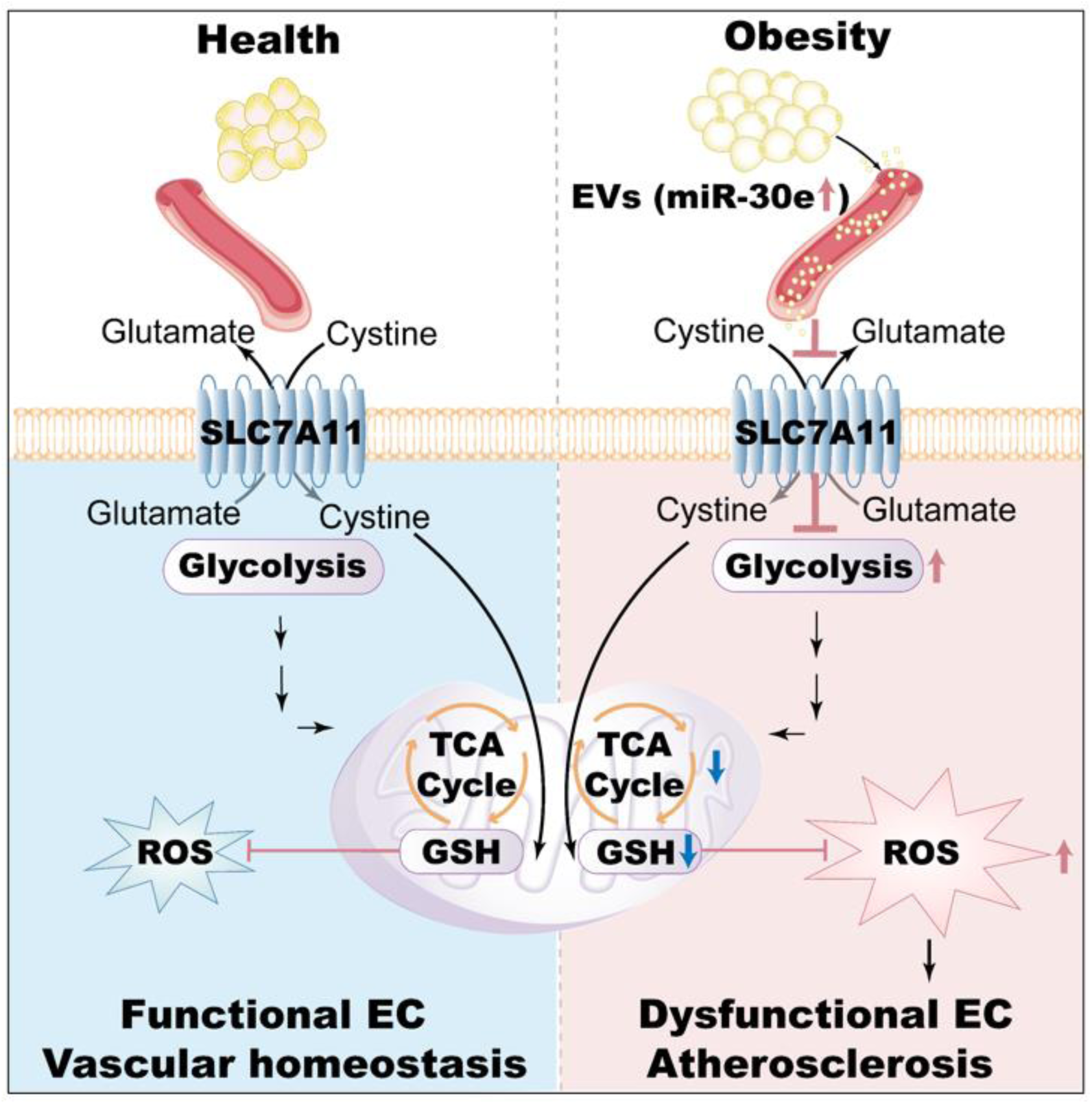
Schematic illustration of adipocyte-derived miR-30e aggravating atherosclerosis via targeting SLC7A11 in vascular endothelium. Under healthy conditions, SLC7A11 in ECs regulates cystine uptake and glutathione biosynthesis, which contribute to redox balance and EC homeostasis. By contrast, obesity increases the adipocyte-derived miR-30e-5p that targets endothelial SLC7A11 in an “endocrine-like” manner. This SLC7A11 downregulation leads to an increase in glycolysis and ROS with a concomitant decrease in oxidative phosphorylation, thus exacerbating EC dysfunction and atherogenesis.

In diet-induced obesity, expanded adipose tissues become deleterious, imposing inflammation and redox stress on the vasculature, thus augmenting atherosclerosis.^9^ Conceptually, adipose tissue constitutes a depot of circulating exosomal miRNAs, a novel mode of cell-to-cell communication regulating gene expression in distant tissues.^12^ Our experiments involving human patients demonstrated a significant elevation of adipose-derived miR-30e-5p in obese individuals with CAD, which was associated with the progression of atherosclerosis. This association was consistent with results from animal experiments revealing increased level of miR-30e-5p in WAT, circulating exosomes, and aortic tissue in obese rodents. In line with our findings, a previous study showed an increase in miR-30e-5p level in circulating EVs in *db/db* mice, which coincided with coronary microvascular dysfunction and rarefaction.^20^ With obesity significantly increasing the risk of cardiovascular impairments, miR-30e-5p may be multi-faceted in the cardiovascular system. Furthermore, our results show that miR-30e-5p–promoted atherosclerosis was independent of LDL-C level, so miR-30e-5p may directly act on the vascular endothelium rather than affecting liver lipid metabolism to regulate atherosclerosis.

Other miR-30e targets that might affect EC functions include adrenoceptor α2A and α2B, phospholipase C γ1 (PLCγ1), and neuronal growth regulator 1 (*Negr1*).^17,34,35^ Data from our snRNA-seq (Figure 3A) indicated that the levels of adrenoceptor α2A and α2B were low in all cell clusters and almost undetectable in ECs. PLCγ1, a direct effector of VEGFR2 and other receptor tyrosine kinases, has an atherogenic effect.^35,36^ However, our snRNA-seq data showed that PLCγ1 level was elevated in adipocytes but decreased in ECs under an HFD. Of note, *Negr1* variants are associated with BMI and obesity.^37,38^ Genetic ablation of *Negr1* in rodents promoted adiposity and increased plasma glucose and insulin levels.^39^ A recent human population study found adiposity and type II diabetes linked to lower serum level of NEGR1.^40^ Consistently, our data revealed that an HFD reduced the level of *Negr1* in both adipocytes and ECs (Supplemental Figure 2C). Obesity can increase other atherosclerotic risk factors, including diabetes, hypertension, dyslipidemia and chronic kidney disease.^41–44^

Providing that NEGR1 has a role in BMI and neuropsychiatric regulation, further studies on miR-30e regulation of NEGR1 in terms of chronic kidney disease syndromes are warranted.

With its enzymatic functions of cystine uptake and GSH biosynthesis, an optimal level of SLC7A11 protects cells against redox and ferroptotic stresses. A previous study showed that the ferroptosis inhibitor ferrostatin-1 reduced iron accumulation and increased the level of Slc7a11 in the aorta of atherosclerotic *Apoe*^-/-^ mice.^24^ Genetic ablation of IL-17D increased the expression of SLC7A11 and decreased atherosclerosis in *Apoe*^-/-^ mice.^45^ Several other miRNAs such as miR-144-3p, miR-181, and miR-429 can also target *SLC7A11* mRNA.^45–47^

Our finding that miR-30e-5p targeting *SLC7A11* mRNA exacerbated atherosclerosis is novel and significant in several ways. First, the reduction of atherosclerosis in *ob/ob* mice administered miR-30e-5p and EC-*Slc7a11*^-/-^ mice recapitulated the causality between obesity-increased miR-30e-5p, EC dysfunction, and atherosclerosis in humans. Second, downregulation of the miR-30e-5p–SLC7A11 axis caused a metabolic shift from glycolysis to mitochondrial oxidative phosphorylation, which enhanced mitochondrial function in ECs. Third, in addition to miR-30e-5p, proinflammatory cytokine TNFα and oscillatory flow also decreased the level of SLC7A11 in ECs (Supplemental Figure 6A). These results suggest that other atheroprone factors synergize with obesity to reduce SLC7A11 level in the vascular endothelium. Because SLC7A11 can suppress ferroptosis, we examined whether miR-30e-5p is involved in ferroptotic regulation. As shown in Supplemental Figure 6B-D, miR-30e-5p overexpression affected neither the level of ferroptosis-related genes such as *GPX4*, *TFRC*, *FSP1* and *ACSL4* nor intracellular Fe^2+^ storage. These results suggest that EC dysfunction caused by the miR-30e-5p–SLC7A11 axis depended on altered central carbon metabolism (i.e., glycolysis and oxidative phosphorylation) and mitochondrial function, rather than the ferroptosis-related mechanism.

The present study has several limitations. Although we established a link between miR-30e-5p and SLC7A11 in the pathogenesis of obesity-related atherosclerosis, the upstream mechanism by which obesity increases the level of miR-30e-5p remains unknown. Previous studies have shown that p53, acting as a transcription factor, can transactivate miR-30e in tumor cells.^48^ Coincidentally, p53 level is increased in WAT of obese rodents and humans.^49,50^ If indeed the miR-30e-5p–SLC7A11 axis is regulated by p53, the precise cell subcluster in adipose tissue where this occurs deserves further elucidation. With the efficacy of the LNP-delivered miR-30e-5p to reduce atherosclerosis, the translational potential of this pre-clinical study requires validation in clinical trials. With other miR-30e targets, the off-target effects associated with the antisense RNA approach also needs to be considered. Nonetheless, this multiomics study elucidates an adipose-vessel crosstalk involving adipocyte-derived miR-30e-5p and its targeted SLC7A11 in vascular ECs. This endocrine-like axis, linking adipose tissue, endothelial dysfunction, and atherosclerosis, may explain in part the ramifications of obesity-caused atherosclerosis.

## Non-standard Abbreviations and Acronyms

Ad-: adenovirus
ASCVD: atherosclerotic cardiovascular disease
BAT: brown adipose tissue
BMI: body mass index
CAD: coronary artery disease
CKM: cardiovascular-kidney-metabolic
EC: endothelial cell
ECAR: extracellular acidification rate
EndoMT: endothelial-to-mesenchymal transition
EVs: extracellular vesicles
eWAT: epididymal white adipose tissue
FACS: fluorescence activated cell sorting
FISH: fluorescence *in situ* hybridization
GSH: glutathione
HAECs: human aortic endothelial cells
HDL-C: high-density lipoprotein cholesterol
HFD: high fat diet
LDL-C: low-density lipoprotein cholesterol
LNPs: lipid nanoparticles
miRISCs: miRNA-induced silencing complexes
NTA: nanoparticle tracking analysis
OCR: oxygen consumption rate
ROC: receiver operating characteristic
ROS: reactive oxygen species
SLC7A11: solute carrier family 7 member 11
snRNA-seq: single-nucleus RNA-sequencing
TC: total cholesterol
TCA: cycle tricarboxylic acid cycle
TG: triglycerides
WAT: white adipose tissue

## Acknowledgements

We thank Drs. Meihe Li, Yuanyuan Wei, Wenbo Yang, Yuanming Xing, Baochang Lai, and Juan Zhou at Xi’an Jiaotong University for their technical assistance and consultation.

## Author Contributions

C.W., J.S., X.Q., Z.-Y.Y., and J.Y.-J.S.: conceived of the study and designed the experiments; C.W., J.S., X.Q., H.W., W.H., X.L., Y.L., X.Z., K.D., Y.L., H.L.,Y.L., S.G., X.Y., T.B., Y.X.: conducted the experiments; C.W., J.S., F.H., X.D.,S.H., and H.L.: conducted *in silico* analysis; C.W., J.S., X.Q., H.W., N.G., J.Z., Z.-Y.Y., and J.Y.-J.S.: analyzed the data and provided the discussion; and C.W., J.S., X.Q., Z.-Y.Y., and J.Y.-J.S.: wrote the manuscript.

## Supplemental Materials

Expanded Materials and Methods Supplemental Table I–VII. Supplemental Figures 1–6.

References^1–7^

